# Hippocampal Engagement during Recall Depends on Memory Content

**DOI:** 10.1101/080309

**Authors:** David A. Ross, Patrick Sadil, D. Merika Wilson, Rosemary A. Cowell

## Abstract

The hippocampus is considered pivotal to recall, allowing retrieval of information not available in the immediate environment. In contrast, neocortex is thought to signal familiarity, and to contribute to recall only when called upon by the hippocampus. However, this view is not compatible with representational accounts of memory, which reject the mapping of cognitive processes onto brain regions. According to representational accounts, the hippocampus is not engaged by recall per se, rather it is engaged whenever hippocampal representations are required. To test whether hippocampus is engaged by recall when hippocampal representations are not required, we used functional imaging and a non-associative recall task, with images (objects, scenes) studied in isolation, and image-patches used as cues. As predicted by a representational account, hippocampal activation increased during recall of scenes – which are known to be processed by hippocampus – but not during recall of objects. Object recall instead engaged neocortical regions known to be involved in object-processing. Further supporting the representational account, effective connectivity analyses revealed that recall was associated with increased information flow out of lateral occipital cortex (object recall) and parahippocampal cortex (scene recall), suggesting that recall-related activation spread from neocortex to hippocampus, not the reverse.

---

Dominant theories of memory hold that the hippocampus (HC) is critical for the retrieval of an episodic memory based upon a partial or associated cue. This form of memory retrieval is often termed recall. Anatomical findings and computational models suggest that hippocampal circuitry is well-suited to such a process (Marr 1971; Teyler and Discenna 1986; McClelland et al. 1995; Rolls 2013). Under this view, recall begins when a cue activates the trace of an associated memory in HC, and is completed when sensory details of the memory are subsequently reinstated in neocortex (Marr 1971; Teyler and Discenna 1986; McClelland et al. 1995; Bosch et al. 2014). Numerous mechanisms have been proposed to explain the privileged role of the HC in recall, including pattern completion (McClelland et al. 1995; Norman and O’Reilly 2003; Rolls 2013), neurogenesis (Aimone et al. 2011) or the construction of sparse, high-dimensional representations (Marr 1971). In contrast, neocortex is assumed to employ different mechanisms, contributing to memory retrieval either by signaling the familiarity of previously encountered items or by providing sensory details when called upon by hippocampus during recall (Miller et al. 1991; Brown and Aggleton 2001; Ranganath 2010; Staresina et al. 2013).

Most theories accounting for the role of medial temporal lobe (MTL) structures in episodic memory retrieval have adopted this mechanistic distinction between HC and neocortex (Teyler and Discenna 1986; Aggleton and Brown 1999; Norman and O’Reilly 2003; Rolls 2010). For example, one influential model has proposed that HC contributes to retrieval – in both recall and recognition tasks – via ‘recollection’, whereas perirhinal cortex (PRC) plays a complementary role in retrieval by providing a familiarity signal (Aggleton and Brown 1999, 2006). In the present article we use 'recollection' to refer to a retrieval mechanism in which partial information is provided as a cue to memory and the missing details are retrieved via a process akin to pattern completion. As such, we take recollection to be the critical mechanism underlying cued recall^1^. Dominant process-based accounts of episodic memory retrieval, such as the account advocated by Aggleton and Brown, imply that the role of each MTL structure in retrieval is constrained by the mnemonic process it underpins (e.g. familiarity versus recollection), regardless of the content of the memory (e.g. single objects versus objects-in-context; Fig.1A). But, more recently, an alternative ‘representational’ view has emerged in which the division of labor within MTL lies not along lines of mnemonic mechanism but along lines of representational content (Fig. 1B).

**Figure 1.**
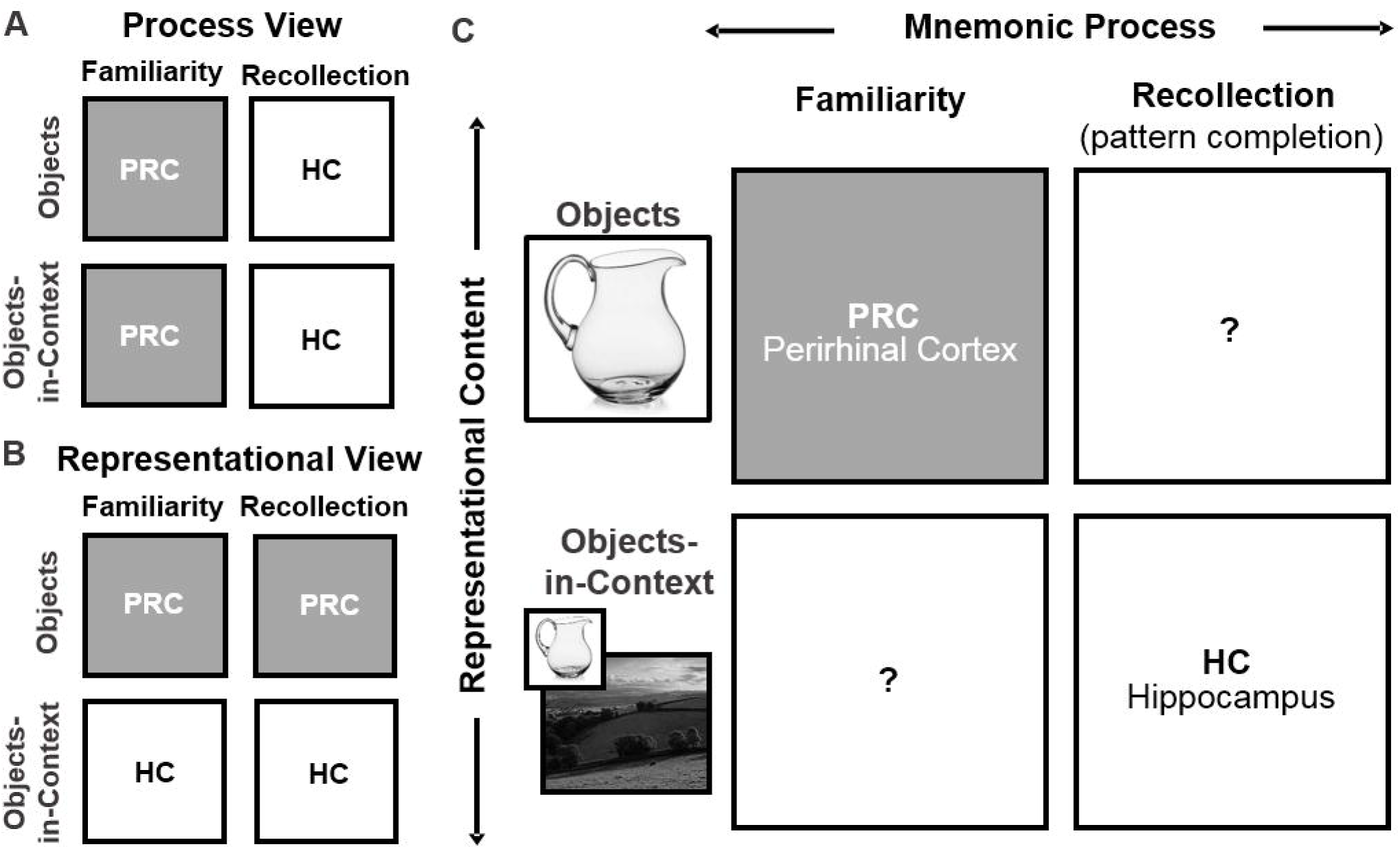
Schematic depiction of theories that explain the role of MTL structures in episodic memory retrieval. The division of labor among MTL subregions may run along lines of (A) mnemonic mechanism, with HC underlying recollection and PRC underlying familiarity, or (B) representational content, with PRC being engaged during retrieval of objects and HC being recruited for retrieval of associative memories, such as an object from a context, or vice versa. (C) A preponderance of the evidence to date comes from tasks that can be solved using item-based familiarity (top left) or via pattern completion of associative memories (bottom right). Very few studies address the complementary scenarios – familiarity for associative memories (bottom left) and item-based recall (top right) – meaning that evidence that can distinguish between the two alternative theoretical accounts is scant. The present study provides the first test of the upper right cell of the matrix by employing a cued recall task, using objects, in which the mnemonic content is non-associative.

The representational view – one version of which is termed the Representational-Hierarchical (R-H) account – rejects the notion that functional distinctions between HC and the surrounding neocortex can be defined in terms of mnemonic processes such as familiarity and recollection (Cowell et al. 2010). Instead, according to representational accounts, functional distinctions are assumed to correspond to differences in the information that each region represents (Bussey and Saksida 2002; Cowell et al. 2010; Graham et al. 2010; Ranganath 2010; Shimamura 2010). Lending support to this theoretical approach, numerous studies employing a range of mnemonic and perceptual tasks have found distinctions in representational content within MTL, showing that perirhinal cortex (PRC) is engaged for processing individual objects, whereas the parahippocampal cortex (PHC) and HC are engaged for processing spatial scenes (Lee et al. 2005, 2008; Barense et al. 2012; Hannula et al. 2013; Mundy et al. 2013; Staresina et al. 2013). In addition, HC appears to play a special role in representing associative relations (e.g. Eichenbaum et al. 1994; Rudy and Sutherland 1995; Diana et al. 2007).

To date, the evidence adjudicating between process-based and representational accounts of episodic memory retrieval remains equivocal (Fig. 1C). A substantial body of evidence implicates PRC in item recognition, which can be supported by familiarity alone (top left cell of Fig. 1C; Meunier et al 1993; Eacott et al. 1994; Winters et al. 2004; Forwood et al. 2005; O’Neil et al. 2009; Schultz et al. 2012), and HC in cued recall of paired associates, which depends on a pattern completion-like process (bottom right cell of Fig. 1C; Hannula et al. 2013; Staresina et al. 2013; Tompary et al. 2016). However, these two findings are in line with both a process view and a representational account. The critical scenarios for differentiating between a process view (Fig. 1A) and a representational view (Fig. 1B) are shown in the bottom left and top right cells of Fig. 1C. These two scenarios, which are less well tested, concern the role of MTL structures in familiarity judgments for associative memories (bottom left cell of Fig. 1C), and in item-based recall (top right cell of Fig. 1C).

According to representational accounts, the contribution of each MTL structure to memory retrieval is determined by its representational content. Thus, the R-H account makes two clear predictions that contrast with a process view. First, it predicts that HC has the capacity to signal familiarity when a memory task employs associative or spatial stimulus material (bottom left cell of Fig. 1B). In fact, there is some evidence for this proposition (Bird et al. 2007; Wais et al. 2010), but this question is not the focus of the present investigation. Instead we focus on a second, more controversial prediction. The R-H account claims that the reason HC is so often implicated in recall (and thereby also in recollection) is that recall is typically performed for associative memories, and associative memories are represented in HC (Cowell et al. 2010). Thus, the second prediction is that the extent of hippocampal engagement during memory retrieval depends on the nature of the stimulus material, not on the requirement to perform recall. If the to-be-retrieved memories do not require hippocampal representations, recall should depend on neocortical regions and may even occur without hippocampal engagement (upper right cell of Fig. 1B; Cowell et al. 2010).

This prediction has not been tested because, to our knowledge, no neuroimaging study has used a retrieval task that decouples the process of recall from the content of the retrieved memory. Neuroimaging studies of recall have almost invariably employed associative material, in which arbitrarily paired images (e.g. an object and scene) are presented at study, and one item is used to cue recall of the other at test (bottom right cell of Fig.1C; e.g. Hannula et al. 2013; Staresina et al. 2013; Tompary et al. 2016). But, such tasks require retrieving information about the association between a pair of distinct images. According to the R-H account, such inter-item associations are optimally represented in HC. This contrasts with, for example, intra-item associations between the parts of an object, which are represented lower down the hierarchy in upstream PRC. Consequently, previous fMRI reports of hippocampal engagement in cued recall (e.g. Hannula et al. 2013; Staresina et al. 2013; Tompary et al. 2016) could be explained either by a process-based account in which HC employs specialized mechanisms for recall, or by a representational account in which HC represents inter-item associations.

To test the prediction that representational content, and not mnemonic process, determines the involvement of MTL regions in recall, we took a standard fMRI recall paradigm and manipulated the stimulus material (Fig. 2A and Supplementary Fig.1). Our critical condition was a non-associative object recall task, in which subjects studied single images of everyday objects and were cued at test with circular patches taken from studied and unstudied images. For the control condition we used an analogous recall task with scene images and scene-patches, because the components of scenes are conjoined by arbitrary associations, for which HC representations are thought critical. Importantly, ‘remember’ responses were associated with explicit recall of the object or scene image and were verified by asking subjects to give the name of the object or scene in a post-scan test (Fig. 2B). We predicted that, although both tasks required retrieval of a whole memory from a partial cue, non-associative object recall would engage HC less than scene recall. Moreover, relative to HC, we expected that neocortical object-processing regions, such as PRC, would be more engaged during object recall.

**Figure 2.**
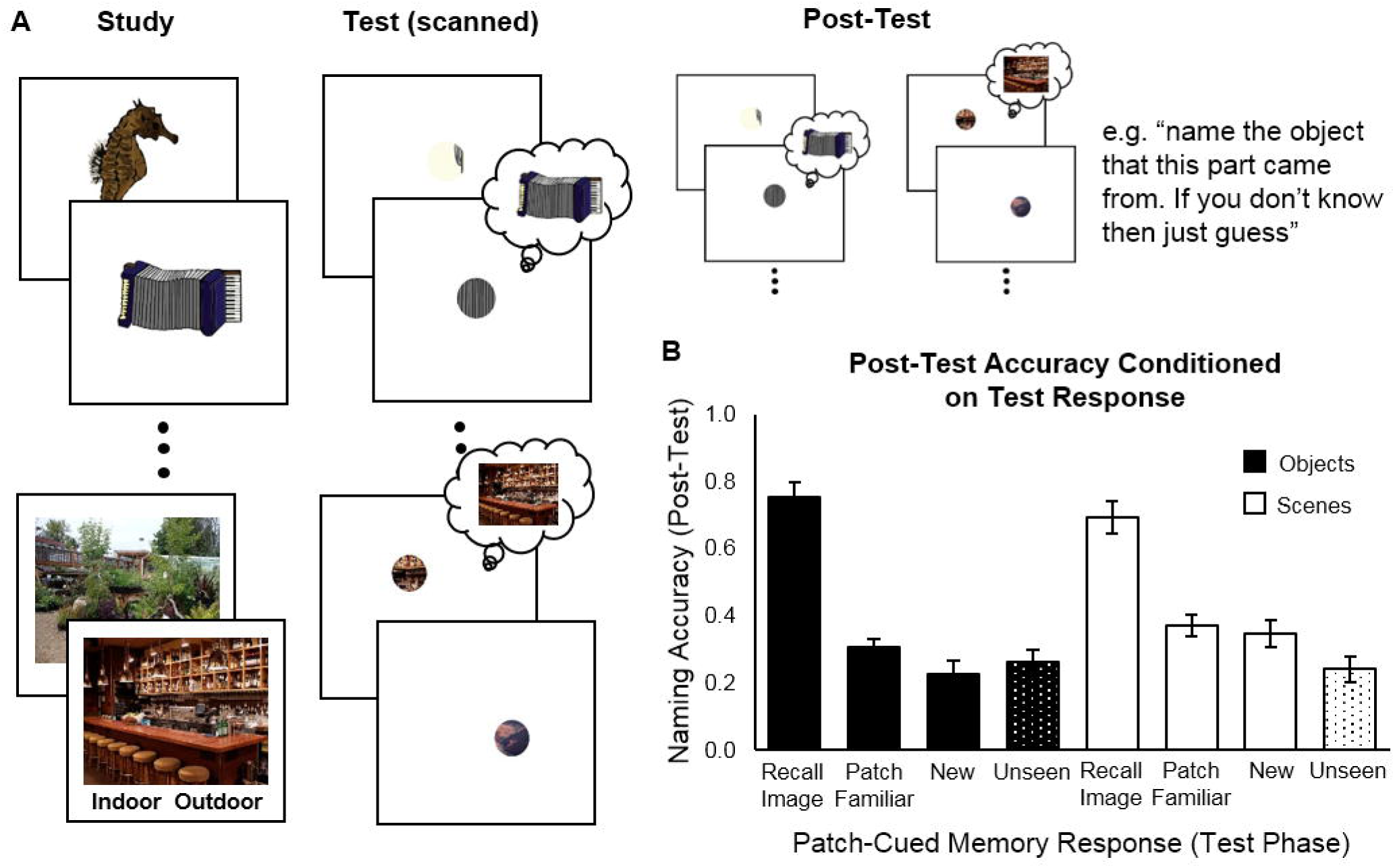
Schematic of task and behavioral results. **(A)** Subjects studied 90 objects and 90 scenes, indicating natural/manmade (objects) or indoor/outdoor (scenes). At test, subjects saw 120 object patches and 120 scene patches (90 seen, 30 unseen) and responded recall, familiar or new, with recall indicating that they specifically remembered the whole image that a patch came from. Finally, subjects saw all 240 image patches a second time and were asked to type the name of the image that the patch came from. If they did not recall seeing the image they were encouraged to guess. **(B)** Proportion of images correctly named at post-test, conditioned on subject response at test. Responses to seen and unseen images are shown separately and the proportion of unseen images correctly named has been collapsed across test response. Error bars show 95% CIs around the means.

In addition to the above analyses, we used Dynamic Causal Modeling (DCM; Friston et al. 2003) to measure effective connectivity between HC, neocortical MTL (PRC, PHC), and an object selective region of lateral occipital cortex (LO). Because DCM permits inferences about the direction of information flow between brain regions, it allowed us to ask which region was driving recall-related neural activity. According to process-based accounts, HC drives the reinstatement of sensory details in neocortex via feedback, but this prediction has been tested only with associative recall tasks (e.g. Staresina et al. 2013; Danker et al. 2016). In contrast, representational accounts predict that recall-related activity will be driven by whichever brain region supports pattern completion of the retrieved memory; importantly, this should be an extra-hippocampal region when the retrieved memory does not comprise associative or relational material. To adjudicate between these accounts, we used DCM to compare models in which recall-related activity was driven by HC with models in which recall-related activity was driven by neocortical sites.

Our univariate results revealed, in line with the predictions of the R-H account, that hippocampal activation was modulated by representational content, increasing during scene recall but not during object recall. In contrast, activation in PRC, which has frequently been implicated in object-processing, did increase during object recall. In addition, our DCM analyses revealed that recall was associated with increased information flow out of neocortical sites for both objects (LO) and scenes (PHC). This suggests that these neocortical regions were driving recall, as predicted by the R-H account, and not merely driven by feedback following successful pattern completion in HC.

## Materials and Methods

### Subjects

Twenty-two (7 male) native English speakers from the University of Massachusetts Amherst and Dartmouth College communities were recruited. Two subjects were excluded from the analyses, one due to excessive motion during scanning, and the other because behavioral responses were not recorded for the majority of their scanned trials. The remaining subjects were between 18 and 32 years of age (M=22.9, SD=3.9) with normal or corrected-to-normal vision. Subjects were paid for their participation and informed consent was obtained in accordance with the University of Massachusetts Amherst and the Dartmouth College Institutional Review Boards.

### Materials

Stimuli were 250 color images, 125 objects and 125 scenes, with 5 images of each type used as practice stimuli (Fig. 2A and Supplementary Fig. 1). Object images were color illustrations (600 × 600 pixels) of natural (e.g. seahorse) and manmade (e.g. accordion) objects taken from a freely available database (Rossion and Pourtois 2004). Scene images were color photographs (600 × 800 pixels) of indoor (e.g. child’s bedroom) and outdoor (e.g. beach volleyball court) scenes, selected to depict non-overlapping contexts (Oliva and Torralba 2001; Quattoni and Torralba 2009; Xiao et al. 2010; Martin Cichy et al. 2014). Cues were circular image-patches, 150 pixels in diameter, taken from various locations on the whole images. Cue patches were chosen to contain minimal semantic information and any other obvious cues that we thought might allow the identity of the whole image to be guessed (i.e. without study). For example, the patch taken from the baby carriage image (object) did not include a whole wheel and the patch taken from the laundromat image (scene) did not include a whole washing machine.

### Procedure and Design

The experimental session comprised an unscanned study-phase (study whole images), a scanned test-phase (patch-cued image recall) and functional localizer (face, house, object, scrambled-object), and an unscanned post-test phase (patch-cued image naming; Fig. 2A). Before starting, subjects completed an abbreviated version of the experimental session – this was primarily so subjects knew that memory responses (remember, familiar, new) made during the scanned test-phase would be verified in a post-test naming task.

### Study phase

The study phase was conducted in a room adjacent to the scanner and functional data were not collected. Subjects viewed 180 images (90 object, 90 scene) sampled from the full set, with the remaining 60 images (30 object, 30 scene) serving as foils at test (across subjects all images served as foils a roughly equal number of times). There were 20 blocks, each comprising nine object or scene images. Blocks of each image type were interleaved and the order was counterbalanced across subjects. Image order was randomized across blocks. Blocks began with a 3 s introduction screen indicating whether the subsequent block would contain object or scene images. Each trial began with a 200 ms fixation, followed by the study image and a text prompt that appeared on screen for 3 s. On object trials subjects indicated whether the object was natural or manmade, and on scene trials subjects indicated whether the scene was indoors or outdoors, making their responses by keypress.

After viewing all 180 images, subjects saw half of the images for a second time with the cue patch removed, effectively creating an aperture through to the background color (see Supplementary Fig.1). As before, subjects responded ‘natural’ or ‘manmade’ for object images, and indoor or outdoor for scene images. The purpose of this manipulation was to increase recall during the test phase (i.e. memory for the whole image) without increasing the familiarity of the image patch. Note, since there were no differences between study-once and study-twice in the imaging data (i.e. there was neither a main effect of study-once versus study-twice nor any interaction involving study condition, all p>.05) we do not discuss this manipulation further in the main text (see Supplementary Fig. 2 and Supplementary Fig. 3).

### Test phase

During the scanned test-phase, subjects were shown all 240 image-patches from the 180 studied images and 60 unstudied images. There were 10 scanned runs, each comprising one block of object cue-patches and one block of scene cue-patches. Within a block there nine of the patches came from studied images and three of the patches came from unstudied images. In addition, there were three baseline trials (Stark and Squire 2001). On baseline trials subjects saw a small fixation point and pressed a button each time it flickered (once or twice), on all other trials a fixation-cross was shown for 250-750 ms (M=500 ms; duration varied randomly on each trial), followed by a cue and text prompt. The cue and prompt remained on screen for 3 s and were replaced by a fixation-cross for 3 s, 5 s or 7 s followed by a 1 s blank screen.

Before entering the scanner, subjects were verbally instructed that they should use the button box to respond ‘remember’, ‘familiar’, or ‘new’ to each image patch.^2^ They were asked to respond quickly but accurately, responding ‘remember’ only if they specifically remembered seeing the whole image during the study phase. In addition, once in the scanner, subjects reviewed text instructions while reference and anatomical scans were completed:

> “You will see part of an object/scene and be asked if you remember what object/scene it came from. The part may be from an object/scene that you have seen before but sometimes the part will be from an object/scene that you have not seen before. You will have three options:
>
> 1. Remember, i.e. you remember the object/scene from the first part of the study.
> 2. Familiar, i.e. you think you saw the part but can’t remember the object/scene it was from.
> 3. New, i.e. you didn’t see the object/scene that the part is from in the first part of the study.”

### Functional localizer

Upon completion of the test-phase subjects completed two passive viewing runs of a functional localizer. Localizer runs consisted of sequentially presented grayscale images of houses, faces, objects and scrambled-objects overlaid with a grid. Each image was presented for 700 ms, with presentation blocked by category. Block order was randomized for the first presentation of each category and this order was subsequently repeated three times for a total of 12 blocks (3 per category).

### Post-test phase

The block structure and trial order were identical to the test phase (with baseline trials omitted). For each patch-cue subjects were prompted to name or describe the corresponding whole object or scene - typing their response. On each trial, a fixation-cross appeared for 200 ms, followed by a cue and text prompt that remained until the subject responded. If a subject did not know the identity of the whole image that a patch-cue came from (i.e. they failed to recall) they were asked to type in a guess.

### Structural and Functional Magnetic Resonance Imaging (MRI)

Imaging was performed using a 3T Philips Achieva Intera scanner with a 32-channel head coil, at Dartmouth College, NH. Each session began with a high-resolution T1-weighted anatomical scan, using an MP-RAGE sequence to acquire 160, 1.0 mm sagittal slices, covering a field of view (FOV) of 240 × 120 × 188 mm (TR, 9.9 ms; TE, 4.6 ms; flip-angle, 8°). Functional scans were acquired using a T2-weighted echo-planar imaging (EPI) protocol (TR, 2000 ms; TE, 30 ms; flip angle, 90°; 3 mm^3^ voxels; matrix size, 80 × 80; FOV, 240 × 105 × 240 mm). Thirty-five axial slices and 142 volumes were acquired per run (141 volumes for one subject), for a total run duration of 284 s.

### Conventional Functional Data Analyses

Functional data were preprocessed and analyzed using BrainVoyager and custom MATLAB code written using the NeuroElf toolbox. T1 scans were registered to the functional scans and the data were interpolated to 1 mm isotropic space and warped to Talairach space (Talairach & Tournoux, 1988). Preprocessing included slice acquisition time correction, 3D motion correction, linear trend removal, temporal high-pass filtering (3 cycles per run) and spatial smoothing using a 6 mm Gaussian kernel.

MTL ROIs were defined in each subject using anatomical landmarks (Pruessner et al. 2000, 2002). In addition, a lateral occipital cortex (LO) ROI was defined by conducting an random effects (RFX) general linear model (GLM) on the functional localizer data, with regressors for the face, house, object and scrambled-object blocks, and placing a sphere (radius 5 mm) on the peak group-level activation following an object minus scrambled-object contrast.

To examine the effects associated with patch-cued object and scene recall, trials were binned by stimulus type (object, scene) and trials corresponding to studied objects or scenes were additionally binned by memory response (recall, familiar, new). Each trial was modeled by a boxcar function beginning at trial onset and ending when the subject made a response – trials on which the subject failed to make a response, or on which the response time was faster than 200 ms or slower than 5000 ms (<2% of trials) were left unmodeled. The boxcar functions were then convolved with a canonical hemodynamic response function (HRF) and entered into a design matrix along with nuisance regressors for motion and a scrub regressors for time points in which the frame displacement exceeded 0.9 (< 5% of time points). Prior to fitting a GLM to the BOLD signal in a given run, the voxel timecourses were normalized by z-transforming the data based on the mean and standard deviation of the baseline segments. We used only baseline segments in calculating the mean and standard deviation because there were different proportions of recall, familiar, and new trials across runs and subjects. Finally, before performing second-level analyses on the parameter estimates from individual subjects, we removed any values that were more than three SDs away from the mean.

### Dynamic Causal Modeling Analyses

We used DCM (Friston et al. 2003) to investigate information flow between neocortex and HC during recall. In DCM, theories about neural dynamics are instantiated in simple neural models (differential equations) that describe the connections between ROIs (i.e. rate of information flow), and the influence of exogenous (e.g. stimulus on, stimulus-off) and endogenous (e.g. recall, no-recall) factors in driving neural activity and modulating connectivity. The neural dynamics predicted by a given neural model are converted into a predicted BOLD timecourse via a hemodynamic model, which has free parameters to allow for variations in the HRF latency and shape between brain regions (Stephan et al. 2004). Finally, after finding parameters that provide the best fit between the observed BOLD timecourse and the predicted BOLD timecourse, it is possible to compare different models. An advantage of this approach (i.e. generating predicted timecourses) is that any comparison of model fit will naturally take account of the uncertainty inherent in the BOLD measure, with both simulations (e.g. Stephan et al. 2008) and neural recording studies in animals (e.g. David et al. 2008) suggesting that DCM is able to uncover neural dynamics that could not be inferred by simply looking at correlations between observed timecourses.

The DCM analyses were conducted in SPM12 using the same preprocessing steps and ROI definitions as were used for the conventional functional analyses. Our neural models (Supplementary Fig. 5) comprised three ROIs - LO, parahippocampal gyrus (PHG; corresponding to PRC for objects and PHC for scenes) and HC – connected to each other by both forward and backward connections (6 connections). Note that we chose to include LO because it is thought to contribute to the recognition of objects whether presented alone or embedded within scenes (MacEvoy and Epstein 2011). The exogenous input into each model was the sequence of studied image trials (i.e. a box car function for each studied image). For half of the models the exogenous input drove the neural response in LO and for the other half it drove the neural response independently in LO and PHG. The choice of LO or both LO and PHG was based on the physiology of the ventral visual stream (Kravitz et al. 2013) and supported by a post-hoc comparison that tested all possible input locations and intrinsic connectivity architectures (Supplementary Fig.6).

The goal of the DCM analysis was to compare models in which recall-related activity was driven by HC to models in which recall-related activity was driven a neocortical ROI. For an ROI to be considered as driving recall, the neural activity within the ROI itself (self-connection) and/or the flow of information out of the ROI (into other ROIs) must *increase during recall*, relative to timepoints in which recall did not occur. To test this, we created three model families, corresponding to the three ROIs that might be driving increased neural activity during recall (e.g. LO, PHG, HC). Within each family there were seven model variations corresponding to all possible combinations of the self-connection and two outward connections (to other ROIs) that could be modulated by recall. Thus, there were a total of 42 models in a model space (i.e. seven models per family x two input locations x three ROIs that could drive recall; Supplementary Fig. 5).

Functional data were concatenated across runs using the spm_fmri_concatenate.m function and separate GLMs were conducted on each subject’s data. Object and scene trials were coded by independent 2x2-factorial design matrices defined by study status (studied, unstudied) x memory response (recall, no recall). The event duration was set at 2 s to allow for sufficient sensitivity (Staresina et al. 2013) and motion parameters were included as nuisance regressors. Functional timecourses were extracted from ROIs in the left and right hemisphere (LO, PHG, HC) by taking the first eigenvariate (similar to the mean timecourse; see Friston et al. 2006) from the 15 voxels with the greatest recall minus no-recall contrast – computed as the difference between the recall/studied beta weights and no-recall/studied beta weights (Stephan et al. 2010; Staresina et al. 2013).

Model fitting was based on maximizing the free energy (Friston et al. 2003), which provides a measure of model evidence that naturally accounts for complexity. The full space of 42 models were fitted separately to the functional data from the left and right hemisphere for both objects and scenes, giving a total of 168 model fits per subject. We then used RFX Bayesian model selection (BMS) to compare the different model families (Penny et al. 2010). In addition, we conducted a fixed-effects (FFX) model comparison to obtain an estimate of the parameters for each DCM and averaged the parameter estimates across the winning family (parameter estimates are reported in Supplementary Table 1).

## Results

### Behavioral Performance

Behavioral data from the scanned test phase confirmed that image patches successfully cued image memory, eliciting a Recall Image response on 36.8% of studied-object trials versus 10.4% of unstudied-object trials (paired t-test, t_19_=10.6, p<.001) and 29.0% of studied-scene trials versus 8.79% of unstudied-scene trials (paired t-test, t_19_=8.52, p<.001). We used Yule’s Q to compute the relationship between study (study, no-study) and recall (recall, no-recall). Yule’s Q is the correlation between pairs of dichotomous variables after accounting for differences in the means (e.g., a bias to respond ‘remember’ would increase studied Recall-Image responses and unstudied Recall-Image responses). Study was strongly correlated with recall response for both objects, Yules Q = .71 (95% CI: .60 – .82), and scenes, Yules Q = .64 (95% CI: .51 – .77), indicating an effect of study on subjects’ tendency to recall images. There was also an effect of study on the judgment of a patch as familiar, assessed by calculating the number of Patch Familiar responses as a proportion of the total number of non-recall (i.e. Patch Familiar and New) responses. Studied-object patches elicited a Patch Familiar response on 43.6% of non-recall trials versus 37.1% for unstudied-objects (paired t-test, t_19_=2.65, *p*=.016). Similarly, studied-scene patches elicited a Patch Familiar response on 50.7% of non-recall trials versus 35.1% for unstudied scenes (paired t-test, t_19_=2.21, *p*=.039). A repeated measures ANOVA indicated that the effect of study (studied, unstudied) on memory response (Recall Image, Patch Familiar, New) was similar for objects and scenes (F_1.54,29.3_=2.85, *p*=.078, 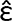=.88). The equivalent analysis of the RT data revealed a main effect of memory response (F_1.42,26_=37.1, *p*<.001, 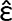=.71)^3^ but no interaction with study condition (studied, unstudied) or stimulus type (object, scene). Recall-image RTs, M=2277 ms, were faster than either Patch Familiar responses, M=2277 ms (t_19_=8.43, p<.001), or New responses, M=2277 ms (t_19_=7.37, p<.001). Patch Familiar RTs and New RTs were not significantly different (t_19_=1.00, *p*=.330; also see Supplementary Fig. 2 and Supplementary Fig. 3).

Finally, to verify that Recall Image responses made during the scanned test phase corresponded to accurate retrieval of whole studied images, we scored the post-test naming responses (Fig. 2B). Naming accuracy was scored by a single rater without knowledge of the study condition (study, no-study) or test-phase response (Recall Image, Patch Familiar, New). Images were not paired with names at study, hence a correct response was somewhat subjective. In line with subject instructions, responses were scored as correct only if they were reasonably specific to the whole image. For example, responding ‘insect’ to the ant image-patch would be marked as incorrect. Conditioning post-test naming accuracy on memory response at test confirmed that naming was more accurate following a Recall Image response than a Patch Familiar response, for objects (paired t-test, t_19_=17.8, *p*<.001) and scenes (paired t-test, t_19_=9.78, *p*<.001). Moreover, recall responses at test (recall, no-recall) were correlated with naming accuracy (correct name, incorrect name) for both objects, Yule’s Q = .80 (95% CI: .73 – .87), and scenes, Yule’s Q = .58 (95% CI: .46 – .70). Note, since naming at post-test is an imperfect measure of memory at the time of test, and scoring was not standardized across objects and scenes, these scores were not used to constrain the functional data analyses. Rather, the purpose of the post-test was to encourage accurate responding at test (subjects knew in advance they would be completing a post-test) and to provide a means of verifying that Recall Image responses in general corresponded to retrieval of whole object or scene images.

### Hippocampal Engagement during Recall Depends on Mnemonic Content

According to the R-H account, the contributions to memory retrieval of subregions within MTL should differ based on representational content (e.g. objects or scenes), not based on mnemonic mechanism (e.g. recall or familiarity). Although it is widely assumed that HC is required for successful recall regardless of mnemonic content, we predicted that the two versions of our patch-cued recall task – objects versus scenes – would differentially engage HC. Specifically, we predicted that patch-cued object recall would elicit much less – if any – HC activation than our control task of patch-cued scene recall. To estimate the contribution of HC and PRC to recall we analyzed only trials in which the cue came from a previously studied image, contrasting trials on which recall was successful (Recall-Image trials) with trials on which it was unsuccessful (Patch-Familiar trials).

We fitted a GLM to the preprocessed functional data (see Materials and Methods) and extracted mean parameter estimates from anatomically defined PRC and HC in each subject. Parameter estimates were submitted to a repeated-measures ANOVA with factors ROI (PRC, HC), image-type (object, scene) and memory response (recall, familiar). A significant 3-way interaction, F(1, 19)=11.65, *p*=.003, confirmed that the engagement of HC and PRC during recall versus familiar trials differed by image type (Fig. 3; also see Supplementary Fig. 4). To investigate the factors driving this 3-way interaction, we conducted separate simple effects ANOVAs within HC and PRC, with factors image-type (object, scene) and memory-response (recall, familiar). Consistent with the predictions of an R-H account, we found a significant image-type x memory-response interaction in HC, F(1, 19)=6.61, *p*=.019, with paired t-tests indicating significantly greater activation during recall-than familiar-responses for scenes, t(19)=4.10, *p*=.001, but not for objects, t(19)=1.78, *p*=.091. Moreover, indicating that the 3-way interaction was driven by the absence of HC activation during object recall, an equivalent repeated-measures ANOVA conducted in PRC failed to find evidence of an image-type x memory-response interaction, F(1, 19)=0.058, *p*=.813, with paired t-tests revealing significantly more activation during recall-than familiar-responses for both objects, t(19)=4.89, *p*<.001, and scenes, t(19)=3.19, *p*=.005.

**Figure 3.**
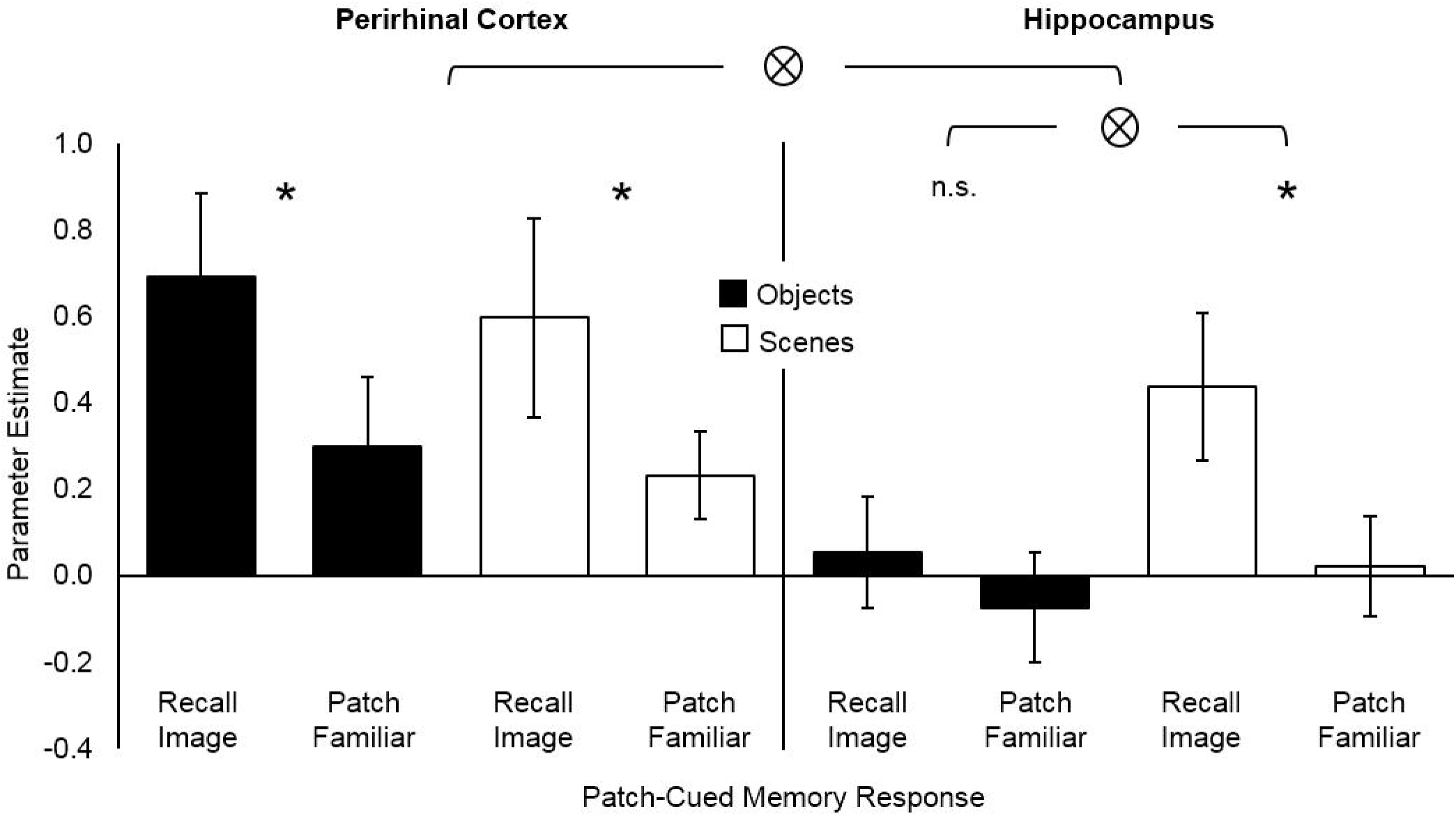
Functional activity during recall and familiar responses to objects and scenes. Parameter estimates shown for perirhinal cortex (PRC; left) and hippocampus (HC; right). Significant (*p*<.05) interactions (⊗) and differences (*) are indicated. Error bars show 95% CIs around the means.

### The Hippocampus at Finer Resolution: Still No Evidence for Engagement by Object Recall

Next, to investigate functional activity in HC more closely, we divided each subject’s HC ROI longitudinally (Fig. 4A) to create three subdivisions corresponding to anterior, middle, and posterior HC (Staresina et al. 2011; Hannula et al. 2013). In line with results from the unitary HC ROIs, a repeated-measures ANOVA with factors subdivision (anterior, middle, posterior), image-type (object, scene) and memory-response (recall, familiar), revealed a significant image-type x memory-response interaction, F(1, 18)=8.35, *p*=.010, but no interaction with subdivision, F(2, 36)=0.014, *p*=.986 (Fig. 4B). Moreover, three simple effects ANOVAs conducted within each HC subdivision separately all revealed a significant interaction of memory-response and image-type (anterior-HC: F(1,19)=4.81, p=.041; middle-HC: F(1,19)=6.00, p=.024; posterior-HC F(1,18)=5.78, p=.027) indicating that hippocampal engagement during recall depends on mnemonic content, even at finer resolution. Finally, paired t-tests conducted separately for objects and scenes in each HC subdivision indicated that in all three subdivisions there was significantly greater activation during recall-than familiar-responses for scenes but not objects: anterior-HC (scenes: t_19_=3.22, *p*=.005; objects: t_19_= 0.920, *p*=.369), middle-HC (scenes: t_19_=4.36, *p*<.001; objects: t_19_= 1.96, *p*=.065), and posterior-HC (scenes: t_19_=3.85, *p*=.001; objects: t_18_= 1.47, *p*=.158).

**Figure 4.**
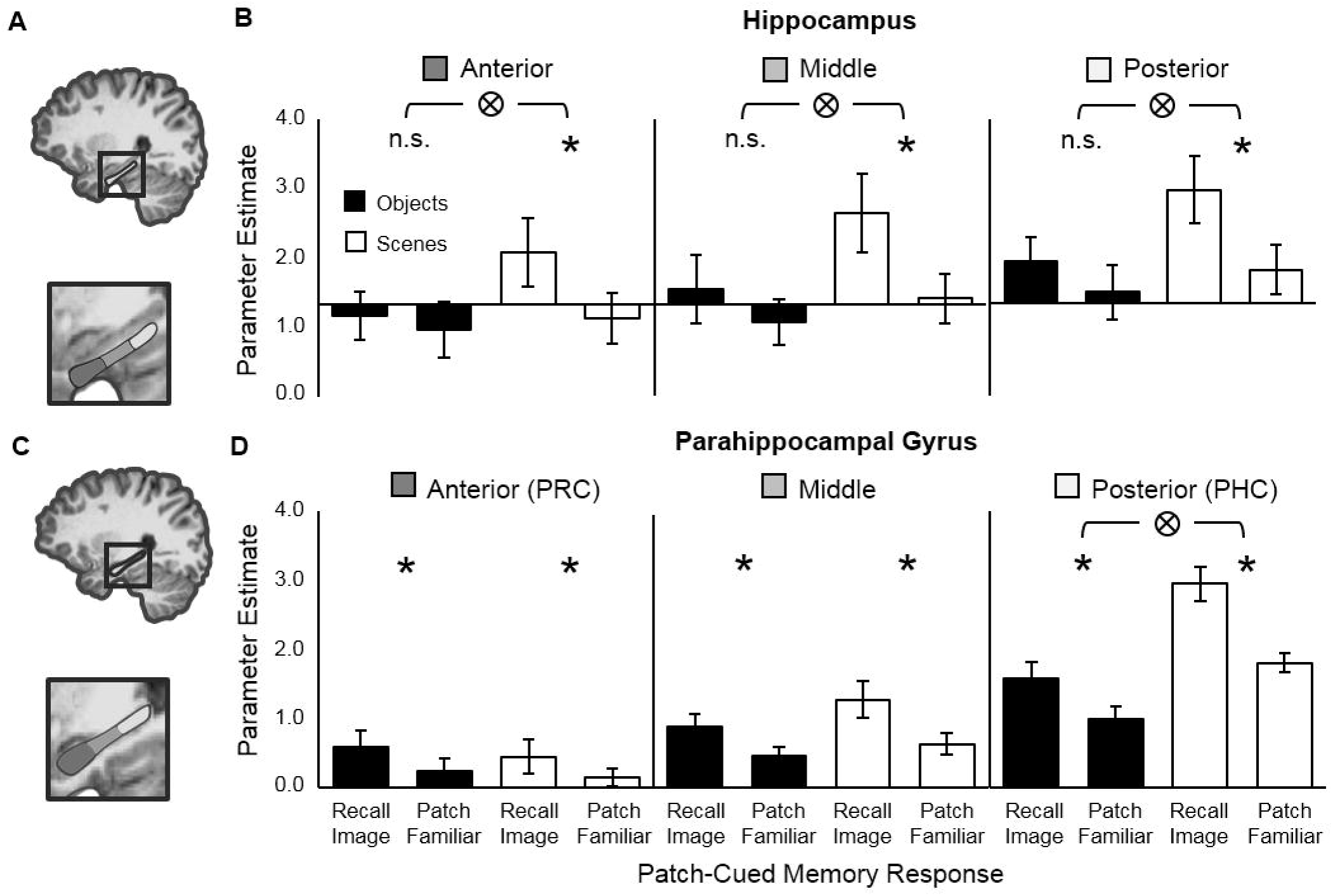
Schematic of 3-part hippocampus (HC) and parahippocampal gyrus (PHG) ROIs and corresponding functional activity from each region. **(A)** Schematic of the 3-part hippocampus ROI. **(B)** Functional activity during recall and familiar responses to objects and scenes in anterior (left), middle (center) and posterior (right) hippocampus. **(C)** Schematic of the 3-part parahippocampal gyrus ROI. **(D)** Functional activity during recall and familiar responses to objects and scenes in anterior (left), middle (center) and posterior (right) parahippocampal gyrus. Note, anterior and posterior parahippocampal gyrus roughly correspond to PRC and PHC respectively. Significant (*p*<.05) interactions (⊗) and differences (*) are indicated. Error bars show 95% CIs around the means.

### The Parahippocampal Gyrus: An Anterior-Posterior Gradient of Engagement by Object and Scene Recall

Prior studies have reported that subregions within the parahippocampal gyrus (PHG) – a neocortical region running parallel to the longitudinal axis of HC and encompassing PRC (equivalent to anterior-PHG) and PHC (equivalent to posterior-PHG) – respond differentially to object and scene recall (Staresina et al. 2011, 2013; Hannula et al. 2013). To look for evidence for this subdivision in our data we defined PHG in each subject using anatomical landmarks, creating three subdivisions along the longitudinal axis, as with HC (Fig.4C). In line with previous studies we expected to find relatively more activation associated with object recall than scene recall in anterior-PHG (corresponding to PRC) than in posterior-PHG (corresponding to PHC). A repeated-measures ANOVA with factors subdivision (anterior, middle, posterior), image-type (object, scene) and memory-response (recall, familiar), revealed a significant 3-way interaction, F(2, 38)=10.38, *p*<.001 (Fig. 4D). An analysis of the simple main effect within each PHG subdivision indicated that there was a significant image-type x memory-response interaction only in posterior-PHG (PHC), F(1, 19)=21.4, *p*<.001, in which – confirming our expectations – significantly more activation was associated with scene than object recall. Finally, to test whether each subdivision of PHG was significantly engaged by recall we conducted paired t-tests separately for objects and scenes, which indicated that in all three subdivisions, for both objects and scenes, there was significantly greater activation associated with recall-than familiar-responses: anterior-PHC (objects: t_19_= 3.56, *p*=.002; scenes: t_19_=2.43, *p*=.025), middle-PHC (objects: t_19_= 5.44, *p*<.001; scenes: t_19_=5.91, p<.001), and posterior-HC (objects: t_19_= 5.28, *p*<.001; scenes: t_19_=11.4, *p*<.001). We note that the asymmetry between PRC and PHC, with more activation during scene than object recall in PHC, but equivalent activation during object and scene recall in PRC, is in line with previous findings (Hannula et al. 2013; Staresina et al. 2013; see Discussion).

### Feedforward Connectivity from Neocortex to HC Increases during Object and Scene Recall

According to process-based accounts, the reinstatement of memory features (i.e. the sensory details of a memory) in neocortex is driven by hippocampal pattern completion (e.g. Staresina et al. 2013; Danker et al. 2016). In contrast, according to the R-H account, the reinstatement of memory features may be driven by pattern completion in any brain region that represents the association between the memory cue and the details that are to be retrieved (e.g. representations of whole objects in the case of our patch-cued object recall task). To adjudicate between process-based and representational accounts (Fig. 5A), the 168 DCMs - 42 models x 2 image types (objects, scenes) x 2 hemispheres (left, right), were individually fit to the observed BOLD timecourses from each ROI (LO, PHG, HC).^4^ The models were split into seven families (Supplementary Fig.5) that differed in terms of the ROI assumed to drive recall (LO, PHG, HC) and the location of the driving input (LO, or LO and PHG) and the evidence in favor of each family evaluated with RFX Bayesian model selection (BMS).

**Figure 5.**
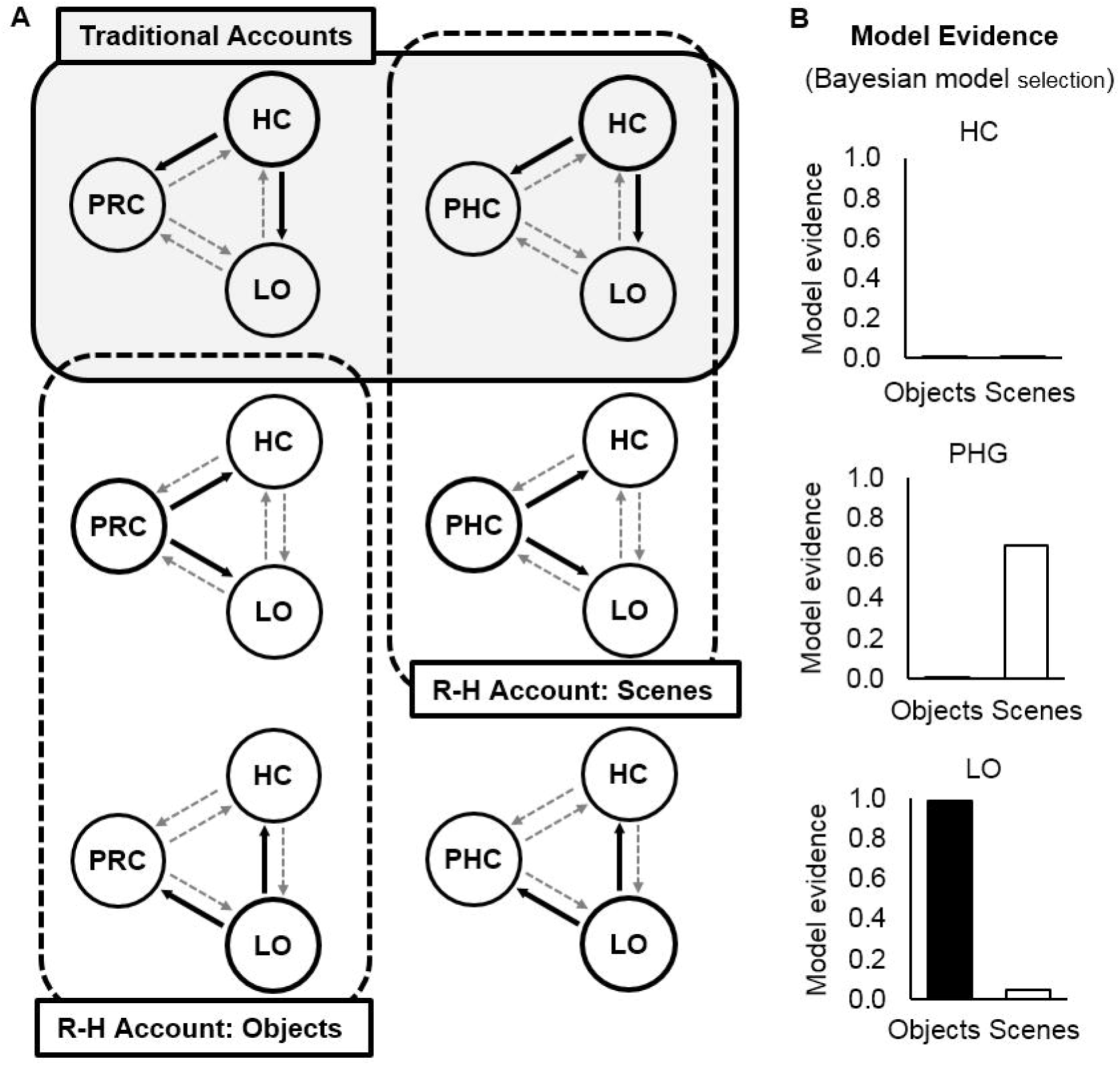
Information flow between ROIs during the cued recall of objects and scenes **(A)** Predictions of traditional and R-H accounts of memory and simplified schematic of the model family space (see Supplementary Fig. 5 for model space). All models had full intrinsic connectivity (grey dashed arrows). Model families were defined by (1) the ROI from which outward connections were permitted to be modulated by recall (bold circles) and (2) the location of the driving input (not shown here). **(B)** Results of the Bayesian model selection for objects (black bars) and scenes (white bars). Results are shown for the winning input family (input into LO for objects, input into LO & PHC for scenes). For objects BMS strongly favored models in which recall modulated information flow out of and within LO (family exceedance probability=.99). For scenes, BMS favored models in which recall modulated information flow out of and within PHC (family exceedance probability=.66). Parameter estimates for modulation by recall are reported in Supplementary Table 1.

Model families were compared by computing the exceedance probability (EP), which is a number between 0 and 1 representing the likelihood of a given model family generating the observed data relative to the likelihoods of the other model families (i.e. the sum of all model EPs is always equal to 1). For objects, BMS strongly favored models with a driving input into LO and recall modulating the connectivity out of and within LO, EP=.99. For scenes, BMS did not overwhelmingly favor any family of model. The winning family had driving input into both LO and PHC, with recall modulating the information flow out of and within PHC, EP=.66. Importantly, even for scenes we found little evidence for families assuming feedback from HC to neocortex during recall (combined EP for families with recall modulating connectivity out of HC was <.001). Looking at individual model EPs (rather than family EPs) confirmed that the most likely model for scenes had inputs into both LO and PHC, with recall modulating information flow out of and within PHC, EP=.70 (Fig. 5B). See Supplementary Materials for parameter estimates (Supplementary Table 1) and supporting post-hoc DCM and PPI analyses (Supplementary Fig. 6 and Supplementary Fig. 7)

## Discussion

We used fMRI and a non-associative recall task to ask whether the engagement of HC during recall is modulated by mnemonic content. This issue is critical for differentiating between a process-based view of memory, in which the functional contribution of distinct MTL structures is best characterized in terms of retrieval processes, and an alternative, representational account of memory, in which the contribution of MTL structures is determined by the content of the memory that is retrieved (Fig.1). As predicted by a representational view, when subjects studied isolated images and were cued to recall them with part of the image, recall-related engagement of HC varied according to mnemonic content. Specifically, HC was engaged during recall of scene images but not during recall of individual objects. In contrast, PRC – known for its role in object processing – was engaged during recall of both objects and scenes. In addition, contrary to the predictions of process-based accounts of memory retrieval, effective connectivity analyses did not support models in which hippocampal feedback drove increased neocortical activation during successful recall. Instead, recall-related changes in effective connectivity were consistent with the R-H account: Bayesian model comparison supported models in which information flow out of LO increased during object recall and information flow out of PHC increased during scene recall. Together, these findings challenge dominant theories of memory retrieval in which HC plays a domain-general role in recall and is engaged regardless of the stimulus material (e.g. Hannula et al. 2013; Staresina et al. 2013; Tompary et al. 2016). Instead, our results suggest that HC does not play a critical role in recall (or recollection) per se, but is engaged to the extent that its representations are required to retrieve a full memory given a partial cue.

While the univariate findings in isolation argue against the simplest interpretation of the process-based view, the DCM results provide important evidence against an alternative formulation of this account. In the simplest version of a process-based account (Fig. 1A), successful episodic memory retrieval is underpinned by activation of a hippocampal memory trace. Under this version of the process-based account, any reinstatement of memory features in neocortex is merely an epiphenomenon of successful pattern completion in HC, and recall should be directly related to increased hippocampal activation. Clearly, this simple version of a process-based view is incompatible with our univariate results, in which recall-related HC activation was contingent upon stimulus content (Fig. 3 and Fig. 4A). However, there is an alternative version of the process-based account, in which HC merely provides an index to memory features in neocortex (e.g. Staresina et al. 2013; Danker et al. 2016). That is, hippocampal pattern completion is necessary but not sufficient for successful episodic memory retrieval; recall also requires the reinstatement of memory features in neocortex. Under this version of the process-based account, the absence of HC activation during recall might simply indicate that pattern completion *per se* elicits negligible activation in HC (because the memory trace itself resides largely in neocortex). But this alternative account nevertheless makes a second, clear prediction: HC is responsible for driving increased activity in neocortex during successful recall. Critically, the DCM analysis revealed virtually no support for models in which HC drove neocortical activation during recall.

Within MTL neocortex, prior work has reported domain-specific contributions to recall (Hannula et al. 2013; Staresina et al. 2013; Vilberg and Davachi 2013), with PRC preferentially engaged by object recall and PHC by recall of scenes. Similarly, we found an interaction between neocortical regions within MTL (PHC, PRC) and stimulus category (object, scene). This interaction was driven by greater recall activation for scenes than objects in PHC, but equivalent recall activation for objects and scenes in PRC. We note that similar asymmetries have been reported previously (Hannula et al. 2013; Staresina et al. 2013). A potential explanation for the involvement of PRC in scene recall is that scenes, and scene parts, often contain objects. Concerning PHC, our findings contrast with prior studies in that PHC was engaged during both object and scene recall – albeit to a lesser extent for objects – whereas previous research has found no engagement of PHC by object stimuli (Buffalo et al. 2006; Hannula et al. 2013; Staresina et al. 2013). One possible reason for the discrepancy is that the BOLD baseline against which we assessed the recall signal differed from previous studies. In Hannula et al. (2013), in the object condition, subjects were shown a scene as a cue and asked to recall the associated object. Thus the BOLD baseline, derived from trials in which recall failed, corresponded to the activation elicited by viewing a complete scene, whereas in our study the BOLD baseline corresponded to the activation elicited by viewing an unrecognizable part of an object. In PHC, one would expect the former baseline condition to elicit higher activation than the latter, which could account for the greater recall effect detected for objects in our study. Critically, there are two key similarities between previous findings concerning neocortical sites within PHG and the present results. First, the interaction between stimulus type (object, scene) and anatomical position within PHG (anterior, posterior, i.e. PRC, PHC) was significant, and second, it exhibited the same asymmetry. We interpret this in terms of a continuum of representations ranging from more object-based in anterior PHG (PRC) to more scene-based in posterior PHG (PHC), superimposed upon an asymmetry wherein objects and scenes come closer to driving neural activation equally in PRC than in PHC.

The literature on episodic memory contains many definitions of a memory retrieval process that requires pattern completion – that is, the filling in of missing details – variously termed recall, recollection or retrieval (as opposed to familiarity; Mandler 1980; Hintzman and Curran 1994; Jenkins et al. 2004; Diana et al. 2007), intentional retrieval (as opposed to automatic retrieval; Jacoby, 1991), or remembering (as opposed to knowing; Tulving, 1985). All of these definitions invoke either conscious awareness or intention, and a subset of them also stipulate that the process retrieves a specific type of information, such as context (Mandler 1980) or autobiographical, episodic details (Tulving 1985b). While definitions of recall or recollection that invoke retrieval of context or personal information undoubtedly apply to episodic recall in many everyday situations, their utility for investigating brain function is limited because they confound a mnemonic mechanism (recollection, or pattern completion) with the content of the memory (e.g. context). In order to discover whether mnemonic mechanism or mnemonic content determines the engagement of different brain regions, we must deconfound the two.

Accordingly, in this study we defined recall as an explicit process that retrieves specific details not present in the environment and confers a feeling of certainty about the past occurrence of an image or event, without stipulating the nature of the retrieved details. In our object recall task, the details to be retrieved were a sufficient number of object features that the object could be identified and prior study of it remembered. Although object-based recall is less commonly invoked than episodic recall as an example of declarative memory retrieval, it provides a critical test for distinguishing the two alternative accounts of the neural basis of cued recall (Fig. 1). Using this test, we demonstrated – contrary to a popular process view – that the contribution of different MTL regions to memory retrieval is better explained by mnemonic content than mnemonic mechanism.

Our patch-cued object recall task has obvious parallels with word-stem completion paradigms. In two studies of amnesic patients, Graf et al. (1984) and Carlesimo et al. (1996) employed word-stem completion under both implicit and explicit task instructions (“complete the stem with the first word that comes to mind” versus “complete the stem with a word from the study phase”), along with both superficial and elaborative processing at the time of study. Our object recall task involved instructions most akin to the ‘explicit instructions’ condition, but encouraged relatively superficial processing by using a super-ordinate categorization task at encoding (objects were judged as “manmade or living”; see Tyler et al. (2004) for evidence that super-ordinate level classification engages earlier visual cortical regions than the cortical regions recruited by basic- or subordinate-level naming). Both Graf et al. (1984) and Carlesimo et al. (1996) reported that amnesic patients – who exhibited declarative memory impairments typically associated with compromised hippocampal function – were impaired at word stem completion only when task instructions were explicit and processing was elaborative. With explicit task instructions but superficial processing, analogous to the present object recall task, amnesic patients were unimpaired relative to controls. Assuming that superficial processing reduces the formation of associations between the studied item and extraneous information, this result is in line with our finding that HC is not engaged for cued retrieval of single objects in a non-associative task. The parallel between our object recall task and stem-completion paradigms thus makes an interesting prediction: HC should be activated during patch-cued object recall if subjects are encouraged to process the objects elaboratively at encoding. In addition, our finding of hippocampal engagement for cued retrieval of scenes can also be compared to the stem-completion results. We suggest that mnemonic content is the critical factor driving the superficial versus elaborative encoding effect in stem-completion. That is, elaborative processing serves to embed single items (words or single objects) into an associative representation, rendering the memory hippocampally dependent. For scenes, however, the encoded information is already relational and does not require elaborative processing to depend on HC. Thus, a second prediction emerges: hippocampal amnesics should be impaired on patch-cued scene recall, regardless of whether they use superficial or elaborative encoding.

One characterization of the differential contributions of MTL structures to memory retrieval is that the PRC represents intra-item associations by performing 'unitization', a process that enables or enhances subsequent familiarity memory for the unitized item (Yonelinas 2002; Mayes et al. 2004; Quamme et al. 2007). The present results are, in part, compatible with this hypothesis: perirhinal unitization of objects at study might facilitate later patch-cued object recall, engaging PRC but not HC at test. However, our task involved recall and not recognition. Only part of an image was presented at test, rather than the whole image. Therefore any unitized, object-level representation could not be activated at test without first retrieving its missing parts. Thus, while perirhinal unitization may play a role in encoding, the contribution of PRC during the retrieval of object memories in our task cannot be one of familiarity signaling alone. Rather, the contribution of PRC is analogous to the role of HC in retrieving episodic memories involving arbitrary associations — a pattern completion process that supplies information not present in the cue. Moreover, the notion of unitization may be redundant, here. A role for unitization in explaining the present results requires that objects, but not scenes, undergo unitization during encoding, rendering only objects hippocampally-independent at test. This explanation thus requires additional assumptions regarding which memories become unitized: is it those items that become hippocampally-independent? The account risks becoming circular. A preferable explanation follows from the R-H account—it is not the process of unitization that is critical to the involvement of HC at the time of retrieval, but the complexity of the conjunctions that the stimuli comprise. Scenes, owing to their complex, relational nature, are represented in HC, whereas objects are not. This uncontroversial assumption is central to the R-H account, and the notion of unitization need not be invoked.

The idea that experience can induce neocortical regions to fill in information via pattern completion has parallels with the perceptual priming literature. It is debated whether the processes and representations underlying visual priming are distinct from, or shared with, those underlying recognition memory (Tulving et al. 1991; Schacter 1992; Berry et al. 2008). A classic demonstration of priming involves the initial inability to comprehend degraded images such as Mooney figures (Mooney 1957) or the famous hidden Dalmatian dog (Gregory 1970), and the ease with which the image contents can be identified after exposure to a coherent version of the image. In such cases, post-priming identification involves cueing the observer with a degraded version of the full image to induce reinstatement of essential information such as global form, and there is evidence that the pattern completion underlying this process occurs in visual cortex (Gorlin et al. 2012). Although our object recall task is directly modeled on standard tests of episodic retrieval (e.g. cued recall of paired associates) it also parallels the part-to-whole completion process required to identify Mooney figures. We suggest that the principal difference between these tasks — Mooney figure priming, object recall and standard recall – is the complexity and specificity of the stimulus representation required for retrieval: representations in visual cortex support superordinate-level categorization based upon Mooney figure cues (Gorlin et al. 2012); representations in PRC underlie retrieval of specific objects in our part-cued object recall task; and hippocampal representations are critical for typical recall tasks in which subjects must retrieve information arbitrarily associated with a test cue. Although the level of explicit awareness associated with retrieval may increase going from Mooney figure priming to fully episodic recall, the retrieval process itself may be extremely similar. Thus, neuroimaging data from both priming and recall tasks may be accounted for by a common pattern completion process acting upon representations at different levels in the ventral visual-perirhinal-hippocampal hierarchy.

Finally, we consider whether the objects in our task were recalled from their part-cues using semantic object knowledge or episodic memory. In line with previous fMRI investigations of episodic memory (e.g. Hannula et al. 2013; Staresina et al. 2013; Danker et al. 2016), we adopted a content-neutral definition of episodic recall, namely, the retrieval of specific details not immediately present, along with certainty about past occurrence, without the requirement that the details be contextual or personal. Our task was tailored to this definition by instructing subjects to give a 'remember' response only when the part cue elicited explicit recollection of the studied image and a high degree of certainty that the image appeared in the study phase. Moreover, post-scan testing verified that subjects identified part cues (e.g. “name or describe the object”) at a much higher rate if they came from objects that were studied (36.8%, where naming rate is collapsed across subject response) than from unstudied objects (10.4%, collapsed across response). This indicates a significant contribution from the episodic memory trace acquired during study to the ability to complete an object from its part. Naming rates were higher still for studied objects that were explicitly recalled in the scanner (75%), but we could not compare these rates to images that were unstudied-but-recalled because such events occurred too infrequently. Both the higher naming rate for recalled images and the infrequency of recall responses to unstudied images suggest that subjects were following instructions regarding 'remember' responses accurately. Thus, the knowledge used to retrieve objects from part-cues was in large part episodic.

Nevertheless, just as semantic knowledge contributes to recall in episodic memory tasks employing more complex, associative information (Bartlett 1932; Bransford and Johnson 1972; Brewer and Treyens 1981) it is highly probable that semantic knowledge contributed in our task. But we do not view this potential contamination as problematic for our interpretation of the present findings. Whereas others (e.g. Tulving 1985b) have tied familiarity-based retrieval mechanisms (“know”) to semantic memory, and recollection responses (“remember”) to episodic memory, we offer an alternative view of declarative memory retrieval. We deliberately avoid one-to-one mappings of process-based distinctions such as familiarity versus recollection onto content-based distinctions such as semantic versus episodic. Instead, we argue for the existence of a continuous hierarchy from early visual regions through PRC into HC, across which a common pattern completion process is employed to retrieve whole representations based upon partial cues. The levels of the hierarchy differ not in terms of process (familiarity versus recollection) but in terms of representational content, with early regions contributing only perceptual information, intermediate regions both perceptual and semantic knowledge, and later (MTL) regions contributing perceptual, semantic and episodic content. A task requiring pattern completion – be it low-level priming, object recall, or episodic recall – will engage regions along this pathway only as far as the representational demands of the task require (Tyler et al. 2004). The contribution of perceptual, semantic or episodic information to retrieval will be determined by representational requirements in the same way.

In sum, our findings suggest that the neural mechanisms underpinning recall do not occur exclusively and mandatorily in HC, and that representational content rather than mnemonic mechanism underlies the functional division of labor within MTL during cued retrieval of episodic memories.

## Author Contributions

Conceptualization, D.A.R. and R.A.C.; Methodology, D.A.R. and R.A.C.; Investigation, D.A.R., P.S. and R.A.C; Data Curation, D.A.R. and D.M.W. Writing – Original Draft, D.A.R. and R.A.C.; Writing – Review & Editing, D.A.R., P.S., D.M.W. and R.A.C; Funding Acquisition and Supervision, R.A.C.

## Acknowledgments

We thank Lila Davachi for helpful discussions, and David Huber, Tim Bussey and Lisa Saksida for discussion and comments on an earlier version of the manuscript. This work was funded by NSF grant 1554871 to R.A.C.

## Footnotes

1 We acknowledge that recollection may also contribute to recognition memory. Brain-based models positing distinct roles of HC versus PRC in episodic retrieval have built upon dual-process models of recognition memory in the cognitive psychology literature (Atkinson and Juola 1973; Mandler 1980; Jacoby 1991; Hintzman and Curran 1994; Yonelinas 1994). But whereas the cognitive dual-process models were originally applied to data from itemrecognition memory tasks, the brain-based models that emerged subsequently have often been applied more broadly, e.g., to item-recognition versus source-recollection (Davachi et al. 2003; Kahn et al. 2004) or to cued recall (Hannula et al. 2013). In this study, we assess the neural basis of recollection in the context of cued recall tasks only; we do not address item recognition memory or associative recognition. Furthermore, our conclusions do not hinge upon confirmation (or refutation) of the dual-process theory of recognition memory

2 We refer to the ‘remember’ response as ‘recall-image’ and the ‘familiar’ response as ‘familiar-patch’ throughout, to avoid confusion between our cued-recall paradigm and recognition memory paradigms. In the present study, at test, subjects were shown only the image-patch, not the whole image. Subjects were instructed to respond ‘remember’ only when they retrieved the whole image from memory. Subjects were instructed to respond ‘familiar’ if the patch seemed familiar without eliciting explicit retrieval of the whole image

3 Six subjects were excluded from this analysis because of missing cells in the unstudied Recall Image condition

4 One sample t-tests performed on the univariate measure of recall activation (i.e. the difference between GLM estimates for Recall Image versus Patch Familiar trials, as described above) confirmed that LO was involved in both object (t_19_=5.03, *p*<.001) and scene (t_19_=5.08, *p*<.001) recall, with no difference in recall activation between image type in LO (t_19_=0.129, p<.899).

